# Hic2 fast-tracks iPS cell generation by suppressing KLF4-dependent epidermal detour

**DOI:** 10.1101/2025.09.11.675597

**Authors:** Meryam Beniazza, Masahito Yoshihara, Daniel F Kaemena, James Ashmore, Suling Zhao, Michael O’Dwyer, Emil Andersson, Victor Olariu, Shintaro Katayama, Abdenour Soufi, Kosuke Yusa, Keisuke Kaji

## Abstract

Reprogramming somatic cells to induced pluripotent stem cells (iPSCs) is the most established cellular conversion by exogenous master transcription factors (TFs). A deeper understanding of this yet inefficient process is critical to extending our capability to control cellular identity for medical applications. Here we report 14 genes essential for efficient iPSC generation, but dispensable for self-renewal. Of those, overexpression of *Hic2*, a transcriptional suppressor highly expressed in PSCs, enhances iPSC generation ∼10-fold. This is achieved through a more direct transition towards pluripotency, bypassing an intermediate state with KLF4-dependent transient epidermal gene expression during iPSC generation. Mechanistically, HIC2 co-occupies these KLF4 targets and directly inhibits their expression. Our work demonstrates that master TFs necessary for cellular conversions can also activate obstructive genes during cellular reprogramming. We propose that identifying transcriptional suppressors against such side effects, like *Hic2*, can be a powerful strategy to achieve more efficient TF-mediated cell conversions.

## Introduction

The successful generation of iPSC with *Oct4, Sox2, Klf4* and *c-Myc* (OSKM) in 2006 demonstrated that cellular identity can be utterly altered by the use of cell type-specific TFs independently from developmental processes^1^. Since then, the use of master TF cocktails became a powerful strategy for the production of desired cell types^2^. Nevertheless, these cell conversions are still inefficient and the resulting cells are often not fully functional. Therefore, as the most well-established TF-mediated cellular reprogramming system, a fundamental understanding of molecular mechanisms underlying iPSC generation is important to further identify ideas and strategies to improve TF-mediated cell conversions.

Generation of iPSCs from fibroblasts requires loss of fibroblastic gene expression and gain of pluripotency gene expression. Accordingly, many genes necessary for the maintenance of a pluripotent state are shown to be also important for pluripotency induction, like *Esrrb*, *Tbx3*, *Utf1* and *Sall4*^3–6^. Loss of somatic cell features, such as a high DNA methylation state and the use of oxidative phosphorylation as a main energy source, is also critical for iPSC generation^7^. In addition to those unidirectional gain or loss of cellular features, iPSC generation involves transient gene expression changes^8,9^. An example is the transient up-regulation of epidermis/keratinocyte genes^8,10^, which has been observed in reprogramming of mouse embryonic fibroblasts (MEFs), B cells and human dermal fibroblasts ^8,10–12^. The epidermis/keratinocyte gene up-regulation is associated with strong exogenous KLF4 expression that is also beneficial to complete faithful reprogramming devoid of abnormal imprinting^10,13^. Moreover, the reprogramming efficiency of keratinocytes is higher than that of fibroblasts^14^. These correlations seemed to suggest a positive contribution of the epidermis-like intermediate state to generate iPSCs. However, two recent single-cell RNA-seq studies investigating the reprogramming process demonstrated that cells in trajectories that do not reach iPSCs tend to have higher epidermal/epithelial genes, indicating that the activation of the epidermal genes constitutes a bottleneck in reprogramming^12,15^.

The identification of genes functionally important during reprogramming informs us of molecular events critical for pluripotency induction. For example, an H3K27 demethylase *Kdm6a* and a Polycomb repressive complex 1 (PRC1) component *Rybp*, have been reported to be important for iPSC generation, while their knockout in ES cells (ESCs) does not affect their self-renewal capacity^16–18^. This has indicated that specific chromatin modification changes play critical roles in this cell fate transition. To further extend our knowledge of essential molecular events during reprogramming, we sought genes necessary for iPSC generation using data from our CRISPR/Cas9-mediated genome-wide knockout (KO) screen^19^. This work led to the identification of 14 genes, not necessary for ESC self-renewal nor proliferation of MEFs, but critical for efficient reprogramming. Among those, overexpression of 7 enhanced iPSC generation, including *Hic2*. HIC2 is a poorly characterised transcription suppressor with a BTB (Broad-Complex, Tramtrack and Bric a brac)/POZ (poxvirus and zinc finger) domain, up-regulated at the late stage of reprogramming. Exogenously overexpressed HIC2 co-occupies KLF4 targets during reprogramming, and directly suppresses transient up-regulation of epidermal genes, allowing the cells to skip the epidermis-like intermediate state, the exit from which usually coincides with a robust up-regulation of pluripotency genes. This work reveals the double-edged sword function of KLF4 in iPSC generation. On one hand, KLF4 is necessary for efficient pluripotency induction. On the other hand, it also induces an epidermal gene expression which precedes and antagonizes pluripotency gene induction. We demonstrate that overexpression of *Hic2* abrogates the detrimental side effect of KLF4 overexpression, rendering reprogramming highly efficient.

## Results

### Identification of novel genes indispensable for efficient pluripotency induction

To deepen our understanding of the molecular mechanisms underlying Yamanaka factors-mediated pluripotency induction, we aimed to discover genes specifically essential for the induction pluripotency but dispensable for its maintenance. To this end, we re-examined the output from the CRISPR/Cas9-mediated genome-wide KO screen performed during reprogramming^19^. Using the MAGeCK software with an FDR<0.15 as a cut-off^20^, we identified a set of 1,298 genes with depleted sgRNAs at the end of the reprogramming process. To pinpoint genes specifically essential for reprogramming, we compared this gene list against two other published genome-wide CRISPR/Cas9 KO screening datasets. The first identified core fitness genes important for general cell survival and/or proliferation^21^, while the second identified genes essential for ESCs’ self-renewal/survival^22^. Of the 1,298 identified genes, 129 genes (9.9% of 1,298 genes) did not overlap with either category (Figure 1A), including *Myc* and *Rybp* which are known to be important for iPSC generation but dispensable for ESC self-renewal^17^. From the 129 candidate genes exclusively essential for iPSC generation, we selected 30 genes with low FDRs in ESC maintenance to validate their unique essentiality during reprogramming (Figure 1B). An interactive platform to view the gRNA depletion FDR comparison between the reprogramming and ESC screens is available at https://kaji-crispr-screen.netlify.app.

**Figure 1.**
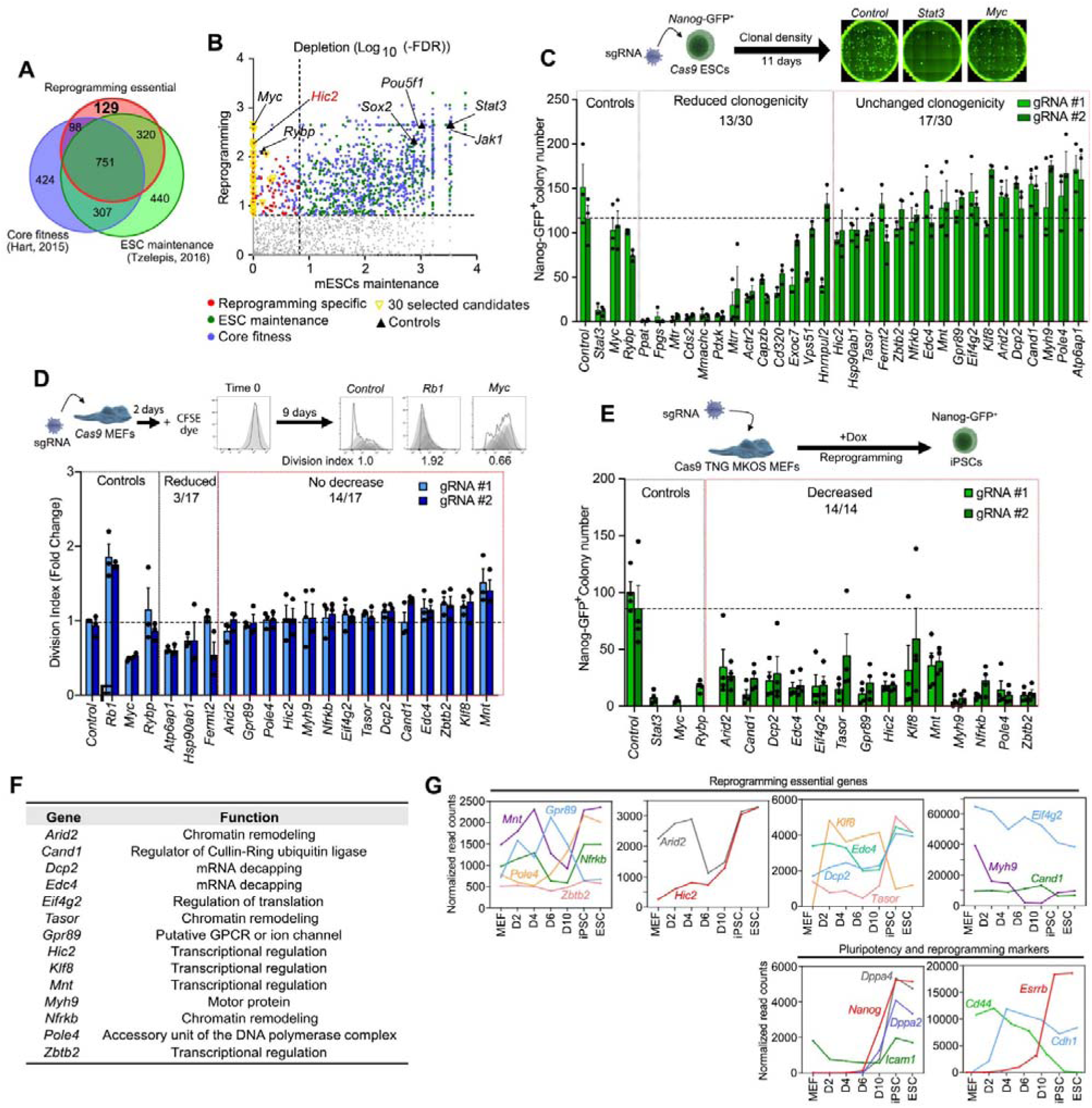
Identification of reprogramming essential genes. **A.** Overlap of 1298 genes with depleted gRNAs in the reprogramming screen with core fitness genes^21^ and ESCs maintenance essential genes^22^. **B.** Comparison of gene depletion FDR in the reprogramming^19^ and the ESC maintenance^22^ screens. Core fitness genes, ESCs maintenance essential genes, and genes specifically important in reprogramming are indicated in blue, green and red, respectively. Triangles indicate 30 genes selected for further validation. **C.** Colony number of Cas9 *Nanog*-GFP^+^ ESCs with sgRNAs expression against the indicated genes. The graph represents an average of 3 independent experiments with 3 technical replicates. Error bars indicate SEM. *P*-values are calculated based on unpaired Student’s t-test and are available in the source data file. **D.** Division index of *Cas9 TNG MKOS* MEFs 9 days after transduction of indicated sgRNAs. The graph represents an average of 3 independent experiments with 2 technical replicates. Error bars indicate SEM. *P*-values are based on unpaired Student’s t-test and are available in the source data file. **E.** *Nanog*-GFP^+^ colony numbers on day 14 of *Cas9 TNG MKOS* MEF reprogramming with sgRNA transduction of the indicated genes. The graph represents an average of 4 independent experiments with 2 technical replicates. Error bars indicate SEM. P-values are based on unpaired Student’s t-test available in the source data file. **F.** List of validated genes specifically important for iPSC generation. **G.** Expression of validated essential genes (top panels), and reprogramming markers (*Icam1, Cdh1, Cd44*), pluripotency markers (*Nanog, Esrrb, Dppa2, Dppa4*) (bottom panels) during reprogramming. The genes were grouped based on their highest expression level.

First, we aimed to validate the non-essentiality of those genes for ESC self-renewal. ESCs with a *Nanog*-GFP reporter and a constitutive *Cas9* expression were transduced with 2 different sgRNAs targeting the 30 candidate genes. The resulting mutant ESCs were cultured at clonal density for 11 days to evaluate their self-renewal capacity (Figure 1C). Out of 30 genes tested, gRNAs against 17 genes did not exhibit a significant reduction of *Nanog*-GFP^+^ colony numbers compared to control sgRNAs (against *Pecam1*) as well as gRNAs against known ESC’s nonessential factors *Myc* and *Rybp*. KO of the other 13 genes showed >50% reduction with at least 1 sgRNA, which were therefore removed from further analysis. Next, we examined if the KO of the remaining 17 affected proliferation of MEFs (Figure 1D). Monitoring the dilution of a preloaded-fluorescence dye by cell division over 9 days, we found that KO of 3 genes moderately decreased MEF proliferation, similar to *Myc* KO. As reprogramming defects caused by KO of these genes might be due to decreased cell proliferation, we excluded these 3 from further examination. Lastly, we confirmed the essentiality of the remaining 14 genes during reprogramming using MEFs with constitutive *Cas9* expression, a *Nanog*-GFP reporter and a doxycycline-inducible *MKOS* cassette (*Cas9 TNG MKOS* MEFs)^19,23^ (Figure 1E). In this reprogramming system, KO of all 14 genes led to a significant reduction in the number of iPSC colonies by at least 50%.

In summary, we have identified 14 genes critical for efficient iPSC generation, but not for ESC self-renewal and MEF proliferation. These genes are involved in various biological processes (Figure 1F), some of which have not been linked with reprogramming before. Interesting examples include *Gpr89*, a putative ion membrane transporter for Golgi acidification^24^, and *Dpc2* and *Edc4* which both encode components of the mRNA decapping complex that cleaves specific subsets of capped mRNA and initiates its decay^25^.

### The addition of Hic2 to the Yamanaka factors leads to a rapid and robust generation of iPSCs

About half of the 14 essential genes we identified are up-regulated during reprogramming and display higher expression levels in ESCs than in MEFs (Figure 1G). This suggests that inefficient up-regulation of these essential genes might represent a bottleneck for reprogramming, and their overexpression might therefore enhance the transition towards pluripotency. We examined this possibility using a *piggyBac* transposon-based reprogramming system (Figure 2A) ^26^. In this assay, overexpression of 7 of the 14 genes enhanced reprogramming efficiency similarly to previously reported reprogramming facilitators, such as constitutively active *Smad3* (*Smad3CA*) and *Rybp* (Figures 2B and 2C) ^17,23^. The most striking effect was obtained upon *Hic2* overexpression, leading to a >10-fold increase in *Nanog*-GFP^+^ colonies over control. *MKOS piggyBac*-mediated reprogramming of neural stem cells (NSCs) was also strongly enhanced by *Hic2* (Figures 2D and 2E), indicating that this phenotype is independent of the somatic cell type of origin. HIC2 (hypermethylated in cancer 2) is best characterized as a transcriptional suppressor, like its paralog HIC1 that interacts with co-repressor CtBP, SIRT1 and HDAC4^27,28^. Endogenous Hic2 expression goes up at the end of reprogramming and peaks in PSCs (Figure 1G), suggesting its importance in facilitating pluripotency gene induction. Because of its remarkable enhancement of iPSC generation, we further investigated the impact of *Hic2* overexpression in reprogramming. We have previously reported that the reprogramming process can be tracked by the expression of cell surface markers CD44, ICAM1, and the *Nanog*-GFP reporter^8^. An analysis of these marker expression changes during reprogramming revealed that *Hic2* overexpression not only boosted reprogramming efficiency, but also dramatically accelerated its kinetics (Figure 2F).

**Figure 2.**
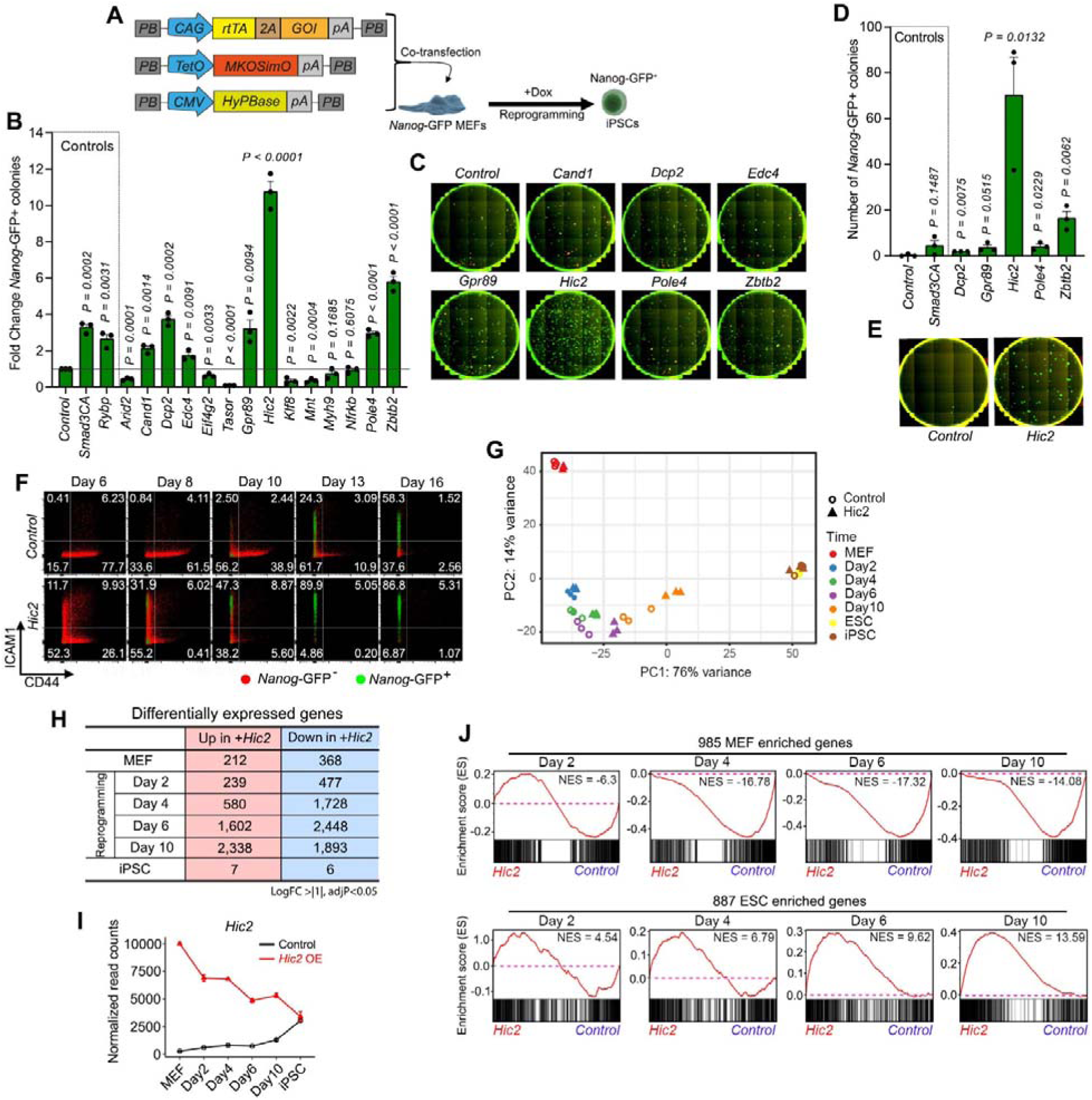
Overexpression of Hic2 alongside Yamanaka factors enhances reprogramming. **A.** *piggyBac* reprogramming with the reprogramming essential genes. GOI; gene of interest. **B.** Fold changes of NANOG^+^ colony numbers obtained after 14 days of *MKOS piggyBac* reprogramming of MEFs with essential genes overexpression (OE) relative to control (Blue Fluorescent Protein, BFP). The graph represents an average of 3 independent experiments, with 2 technical replicates. Error bars indicate SEM. P-values are calculated based on unpaired Student’s t-test. **C**. Representative images of wells with statistically significant reprogramming enhancement in **A**. Red; mOrange, Green; NANOG immunofluorescence. **D**. Number of Nanog-GFP^+^ colonies at day 16 after *MKOS piggyBac* reprogramming of NSCs with a selection of essential genes or BFP (Control) OE. The graph represents an average of 3 independent experiments, with 2 technical replicates. Error bars indicate SEM. P-values are calculated based on unpaired Student’s t-test. **E**. Whole-well images of NSC reprogramming with BFP (control) and Hic2 OE. Green; NANOG immunofluorescence. **F**. Representative CD44/ICAM/*Nanog*-GFP expression changes during reprograming with OE of Hic2 (n=3). Red; Nanog-GFP^−^ cells, Green; Nanog-GFP^+^ cells. **G.** PCA of bulk RNA-seq data from reprogramming time-course experiments in the presence of Hic2 or BFP (control) overexpression. **H.** The number of differentially expressed genes. **I.** Expression of Hic2 during reprogramming with Hic2 or BFP (control) OE. Error bars indicate SEM from 3 replicates. **J.** GSEA with 985 MEF (top) and 887 ESC (bottom) enriched genes during reprogramming in the presence of Hic2 or BFP (control) OE.

Bulk RNA-seq at different time points of reprogramming demonstrated that *Hic2* overexpressing cells start becoming closer to ESCs/iPSCs compared to their cognate control cells as early as day 4 in principal component analysis (PCA) (Figure 2G). Importantly, iPSCs generated in the presence of exogenous Hic2 were highly similar to control iPSCs with negligible differences in gene expression (a total of 13 differentially expressed genes (DEGs)) (Figure 2H and Supplementary Table S1), accompanied by down-regulation of exogenous *Hic2* expression in iPSCs (Figure 2I). Gene set enrichment analysis (GSEA) with gene sets consisting of 985 MEF or 887 ESC enriched genes (MEF vs ESC, FDR<0.05, FC>5) showed significantly accelerated MEF gene silencing in *Hic2* overexpressing cells from day 4, followed by accelerated ESC gene up-regulation at day 6 compared to control cells (Figure 2J)^29^. The total number of down-regulated genes were also larger than that of up-regulated genes in *Hic2* overexpressing MEFs as well as at day 2, 4, and 6 of reprogramming (Figure 2H). These results indicate that HIC2 predominantly acts as a transcriptional suppressor in this context, consistent with previously reported HIC2 function^27^, and this activity could be the main mechanism behind accelerated reprogramming.

### Hic2 fast-tracks iPS cell generation bypassing an epidermis-like state

We further analysed the Hic2-mediated reprogramming acceleration using the *piggyBac* reprogramming system and 10X Genomics’ single-cell RNA sequencing (scRNA-seq) technology (Figure 3A, data available at https://kkaji.shinyapps.io/230420shiny/?_ga=2.71125584.1988667254.1682608652-2076402475.1682608652.). We captured a total of 13,315 cells from MEF, days 2, 4, 6, 9 and 12 of reprogramming and ESCs (Supplementary Figures 1A and 1B). When the cells were plotted in a two-dimensional force-directed layout (FDL)^15,30^, it became apparent that Hic2 overexpressing cells take a much shorter and more direct trajectory towards the pluripotent state, compared to cells undergoing normal *MKOS* reprogramming (Figure 3B and Supplementary Figure 1C). Previously Schiebinger, et al. has analysed the MEF reprogramming process, and in which the cells with trophoblast gene expression are the population closest to the final fully iPSC state^15^. Overlaying this and our data revealed that the cells approaching to a pluripotent state in Hic2 OE reprogramming do not overlap with this trophoblast-like population (Supplementary Figure 1D). Then, we divided the cells with similar gene expression into 12 clusters using Seurat for further analyses (Figure 3C) ^31^. MEFs mainly fall in cluster 1, with minor populations in clusters 10, 11, and 12, indicating heterogeneity of MEFs (Figure 3D). The majority of ESCs belong to cluster 9. Clusters 2, 3, 4, and 5 on the long route toward the pluripotent state are composed of cells undergoing control MKOS reprogramming (Red colours, Figure 3D). Clusters 6, 7 and over 70% of cluster 8, which bridges late reprogramming intermediates and ESCs, are composed of Hic2 overexpression samples (blue colours, Figure 3D). This confirmed that exogenous *Hic2* expression facilitated a distinct and smoother transition to the pluripotent state.

**Figure 3.**
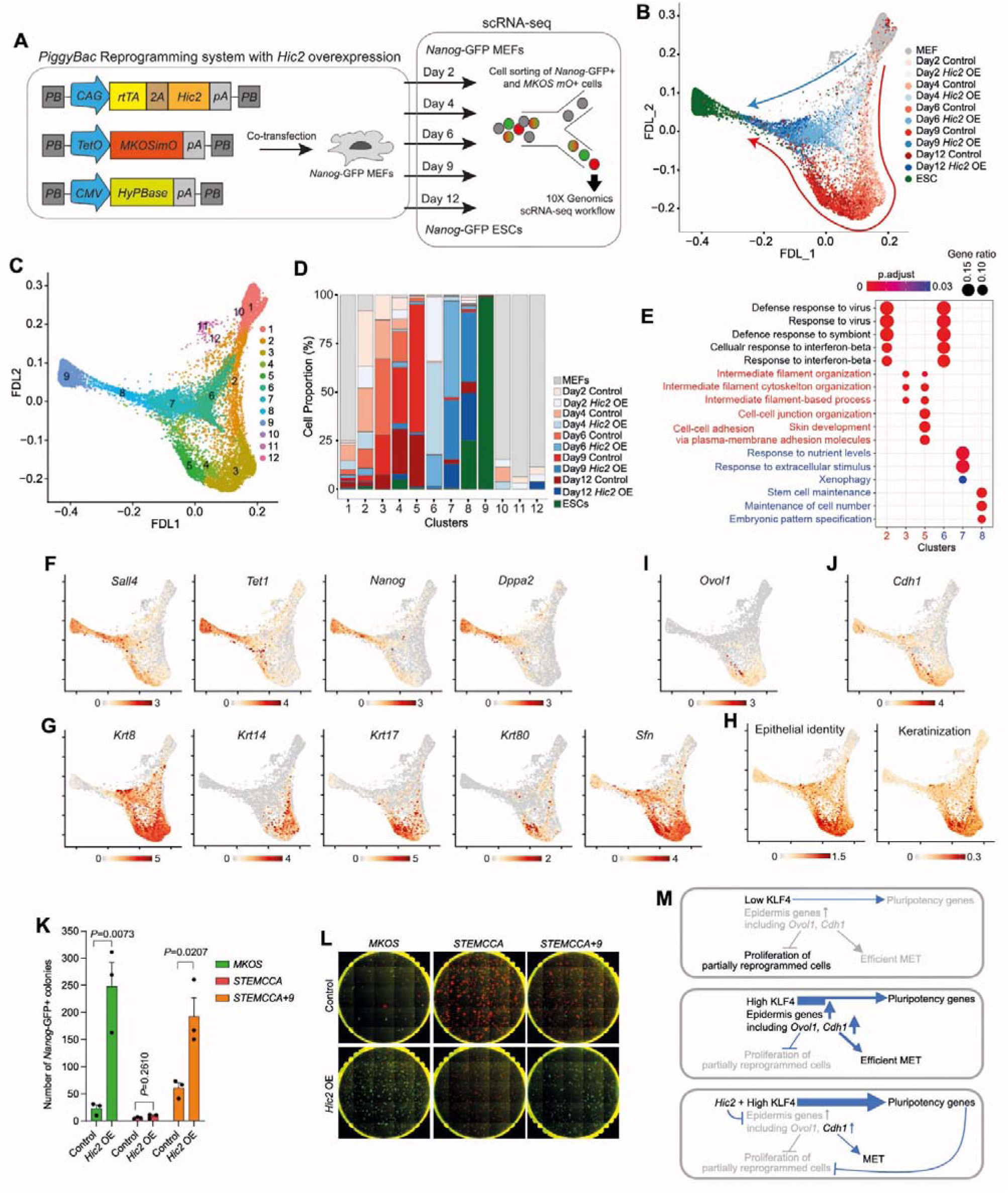
Hic2 fast-tracks iPS cell generation bypassing an epidermal gene expressing state. **A.** Schematic diagram of *piggyBac* reprogramming with Hic2 overexpression and scRNA-seq experimental design. **B.** A two-dimensional force-directed layout (FDL) with sample identity. Grey: MEFs, Green: ESCs, Blue: Hic2 overexpressing cells, Red: control cells. The blue and Red arrows indicate “short” and “long” routes to pluripotency, respectively. **C.** Clustering the cells into 12 groups by Seurat FindNeighbours function. **D.** Proportions of cells that belong to each cluster. **E.** GO terms enriched in the marker genes of the long (2-5) and short (6-8) trajectory clusters. **F-H.** Expression of pluripotency (F), epidermal (G), epithelial identity and keratinization genes (H). **I, J.** Expression of KLF4 targets *Ovol1* (I) and *Cdh1* (J). **K.** Numbers of NANOG+ colonies 14 days after *piggyBac* reprogramming with *MKOS*, StemCCA or StemCCA+9 cassettes in the presence of *Hic2* overexpression or a control (*BFP*). The graph represents an average of 3 independent experiments, with 2 technical replicates. Error bars indicate SEM. P-values are calculated based on unpaired Student’s t-test. **L.** Representative images of I. Red; mOrange, Green; *Nanog*-GFP. **M.** Distinct molecular events in reprogramming with low (top) and high (middle) KLF4 expression or high KLF4 expression with *Hic2* overexpression (bottom).

Next, we performed gene ontology (GO) enrichment analysis using marker genes from the long (2, 3, 4 and 5) and short (6, 7 and 8) trajectory clusters (adjP < 0.05, pct.1/pct.2 > 2) to identify features of each cluster (Supplementary Table S2). The top 3 most enriched GO terms from each cluster are listed in Figure 3E, except cluster 4 which is not enriched in any GO terms. GO term “Stem cell maintenance” is enriched in cluster 8 as expected, with marker genes *Sall4, Tet1, Nanog* and *Dppa2* (Figures 3E and 3F). GO terms related to the intermediate filament and skin development are enriched only in the long trajectory clusters 3 and 5, owing to the expression of multiple keratin genes and other genes highly expressed in keratinocytes, like *Sfn* (Figure 3E and 3G). We and others previously reported transient up-regulation of epidermis/epithelial/keratinocyte genes as a hallmark of cells undergoing Yamanaka factors-mediated reprogramming^8,10–12^. Examination of defined gene sets for epithelial identity and keratinisation also confirmed the epidermis/keratinocyte-like state in the long reprogramming trajectory^15^ (Figure 3H). Violin plots also confirmed homogenous suppression of those genes by *Hic2* at days 4, 6, 9 and 12 of reprogramming (Supplementary Figures 1E and 1F). Recently, Kagawa, *et al.* revealed that transient up-regulation of the epidermal genes during reprogramming are less prominent when reprogramming is carried out with low KLF4 expression. In such reprogramming condition, profound proliferation of partially reprogrammed cells and inefficient mesenchymal-epithelial transition (MET) and pluripotent gene up-regulation is particularly evident^10^. When we examined genes up-regulated in a high KLF4-dependent manner during reprogramming^10^, 78% of detected genes in our scRNA-seq data were enriched in the long reprogramming trajectory (Supplementary Figure 1G). This includes epidermal transcription factor *Ovol1*, which is responsible for the suppression of partially reprogrammed cell proliferation (Figure 3I)^10^. Of note, *Cdh1* (*E-Cadherin*) is a hallmark of mesenchymal-epithelial transmission (MET), but its suppression by *Hic2* overexpression was not as evident as that of other epidermal genes, consistent with the fact that *Cdh1* expression during reprogramming is not transient (Figures 3J).

The circumvention of the transient epidermis-like state in the short reprogramming trajectory prompted us to examine the impact of Hic2 overexpression in a reprogramming context where little epidermal gene up-regulation can be observed. To this end, we used a polycistronic reprogramming cassette referred to as STEMCCA, known to express a low level of KLF4^10,32–34^. This reprogramming cassette generates many colonies, but most of them fail to up-regulate the *Nanog*-GFP reporter, unlike the *MKOS* cassette which expresses a higher level of KLF4 (Figure 3K and 3L). Overexpression of *Hic2* in this context did not increase the number of *Nanog*-GFP^+^ iPSC colonies at all, while the size of partially reprogrammed colonises seemed to become smaller (Figures 3K and 3L). Kim, *et al.,* have reported that *Klf4* cDNA in the STEMCCA reprogramming cassette lacks a sequence encoding the first 9 amino acids, and supplementing the missing 9 amino acids (+9) increases the stability and expression level of KLF4^33^. The resulting STEMCCA+9 cassette increased the number of *Nanog*-GFP+ colonies compared to the STEMCCA cassette (Figures 3K and 3L). When exogenous *Hic2* was supplemented to the STEMCCA+9 reprogramming, we observed remarkable reprogramming enhancement, similar to what is seen with the *MKOS* cassette (Figures 3K and 3L), demonstrating that *Hic2*-mediated reprogramming enhancement is KLF4 expression level dependent.

These data indicate that *Hic2* overexpression facilitates the induction of pluripotency, at least in part, via the suppression of transient epidermal gene up-regulation, i.e. blocking the acquisition of an alternative cellular identity, mediated by high KLF4 expression. Reprogramming with low KLF4 leads to poor induction of epidermal genes, including *Ovol1* and *Cdh1*, resulting in the proliferation of partially reprogrammed cells, inefficient MET and pluripotency gene inductions (Figure 3M, top). On the contrary, reprogramming with high KLF4 induces KLF4-target epidermal genes including *Ovol1* and *Cdh1*, leading to suppression of partially reprogrammed cell proliferation and more efficient MET and pluripotency gene induction (Figure 3M, middle). Nevertheless, only cells that successfully down-regulate the epidermal genes can reach a fully reprogrammed state. *Hic2* overexpression prevents the induction of the epidermal cell identity induced by KLF4^35^, a conflicting cell fate choice, and facilitates the acquisition of pluripotency (Figure 3M, bottom).

### HIC2 co-occupies KLF4 target genes and directly suppresses epidermis gene activation during reprogramming

How does HIC2 prevent KLF4-mediated epidermal gene up-regulation? We initially speculated that *Hic2* overexpression alters KLF4 binding pattern during reprogramming. However, this hypothesis was dismissed as KLF4 binding at 48 hrs of reprograming with and without *Hic2* overexpression was very similar in terms of both location and intensity (Figures 4A and 4B). Instead, unexpectedly, we observed a considerable overlap between HIC2 and KLF4 binding loci in *Hic2* overexpressing reprogramming (Figure 4C), including at the epidermal gene loci suppressed by *Hic2* (Figure 4D and 4E). Of the 45,047 KLF4 ChIP-seq peak loci, 26,876 (∼60%) overlapped with HIC2 ChIP-seq peaks, and their peak summits were located only within a dozen base pairs (bp) away from one another (Figure 4F). In fact, the most enriched TF motif in HIC2 peaks was the KLF1 motif (Figure 4G). In order to examine the impact of HIC2 binding on KLF4 target gene expression, we performed k-means clustering of differentially expressed genes (DEGs) of *Hic2* overexpressing versus control reprogramming (adjP <0.05 and log2FC>|1|), focusing on genes with HIC2 and KLF4 binding within +/-5 kb from the transcription start sites (TSS) (Figure 4H). Of the 5 clusters, scaled expression of clusters 1, 2, and 3 showed suppression by *Hic2*, which is evident as early as day 2. Consistently with the scRNA-seq data, cluster 3 with the transient up-regulation pattern is enriched in “epidermal development” and “keratinocyte differentiation” genes (Supplementary Figure 2). In contrast, genes in clusters 4 and 5 showed higher expression in *Hic2* overexpressing samples after day 4 of reprogramming. Genes belonging to these clusters are highly expressed in iPSCs, including genes with GO terms “stem cell population maintenance” in cluster 5 (Supplementary Figure 2). Thus, this enhanced gene up-regulation could be a consequence of accelerated reprogramming by *Hic2* overexpression. Finally, we confirmed that overexpression of *Klf4* alone is sufficient to induce the expression of epidermal genes in MEFs (Figure 4I). This effect is suppressed when *Hic2* is co-expressed (Figure 4I), demonstrating the direct regulation of epidermal genes by KLF4 and HIC2.

**Figure 4.**
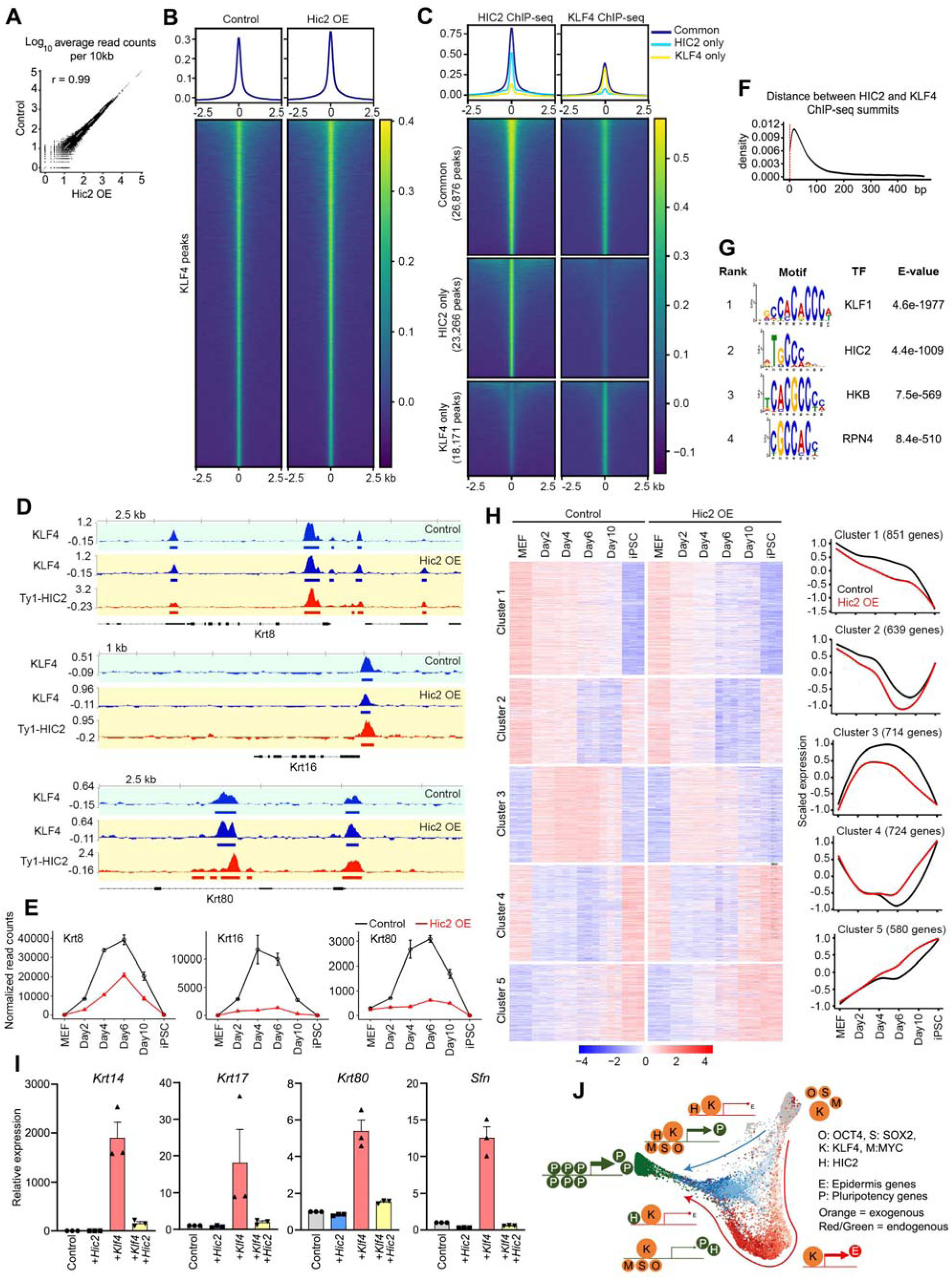
HIC2 co-occupies KLF4 binding sites and suppresses transient gene expression during reprogramming. **A.** Correlation plot of KLF4 ChIP-seq signals in control *BFP* or *Hic2* overexpressing cells after 48 hours of *MKOS* expression. **B.** Heatmaps of the KLF4 ChIP-seq peak regions in control *BFP* or *Hic2* overexpressing cells after 48 hours of MKOS expression. **C.** Heatmaps of the common (top), HIC2-specific (middle), and KLF4-specific (bottom) ChIP-seq peak regions after 48 hours of MKOS expression in the presence of *Hic2* overexpression. **D.** KLF4 (blue) and HIC2 (red) binding at epidermal gene loci in control (top) and *Hic2* overexpressing cells (middle and bottom) at 48 hours of reprogramming. **E.** Expression of the epidermal genes in D during reprogramming with and without *Hic2* overexpression. Error bars indicate SEM from 3 replicates. **F.** Distance between HIC2 and KLF4 ChIP-seq peak summits in the common peaks. **G.** Motif enrichment analysis of HIC2 ChIP-seq peaks by MEME-ChIP tool. H. k-means clustering of differentially expressed genes between control and Hic2 overexpressing reprogramming with KLF4 and HIC2 binding within 5 kb of transcription start sites (left), and scaled expression of genes belonging to each cluster (right). I. RT-qPCR for epidermal genes 96 hrs after overexpression of Hic2, Klf4 or Hic2 plus Klf4 in MEFs. Data represent an average of 3 independent experiments with 3 technical replicates. Error bars indicate SEM. P-values are calculated based on unpaired Student’s t-test. J. A model of Hic2-mediated acceleration of iPSC generation in the presence of robust KLF4 expression.

In summary, these data demonstrate that HIC2 co-occupies a large proportion of KLF4 target loci during reprogramming, and directly suppresses the transient acquisition of KLF4-mediated epidermis-like state, skipping which leads to an accelerated transition to the pluripotent state.

## Discussion

OSKM-mediated reprogramming is a highly complex process that involves multiple stages and events beyond the mere silencing of somatic genes and the activation of the pluripotency gene network. Our study aimed at expanding our knowledge of genes specifically involved in the transition from a somatic to a pluripotent cell state. Of the 14 reprogramming process-specific essential genes we identified, overexpression of *Hic2* drastically enhanced reprogramming efficiency. We showed that this is accompanied by the suppression of epidermal genes normally induced by a high level of KLF4 expression^8,10^. *Cdh1* is also one of the genes efficiently induced by high levels of KLF4^10^. However, suppression of *Cdh1* induction by *Hic2* was modest (Figure 3I and Supplementary Figure 3A). This could be because *Cdh1* is also a direct target of OCT4 and SOX2 in ESCs (Supplementary Figure 3B), where many of the epidermis genes have little OCT4 and SOX2 binding (Supplementary Figure 3C). Reprogramming with high KLF4 expression is beneficial for efficient pluripotency gene induction. Nevertheless, it also results in the epidermis gene activation (Figure 4J)^35^, probably due to the chromatin states and/or lack of suppressors. This leads the cells to transit through an alternative state from which only a fraction of them reach pluripotency. In fact, scRNA-seq analysis during OSK-mediated reprogramming by Guo *et al.* identified a refractory cell population which expresses many high KLF4 target genes, particularly epidermal genes (Guo et al., 2019), features that are also shared with partially reprogrammed iPSCs^36,37^. In reprogramming with retrovirus vectors, transgene silencing has been shown to correlate with a fully reprogrammed pluripotent state, consistent with that pluripotent cells have a high *de novo* DNA methylation activity necessary for efficient viral vector silencing^38^. Accumulating data suggest that down-regulation of exogenous KLF4 at the right timing is rather a necessary step to suppress epidermal genes and complete the acquisition of the pluripotency. Endogenous *Hic2*, a transcription suppressor essential for efficient iPSC generation, is also a target of OCT4, SOX2 and KLF4 in ESCs (Supplementary Figure 3D). Its up-regulation correlates with down-regulation of epidermal genes (Figure 3G, Supplementary Figures 3E and 3F), which suggests that endogenous HIC2, similarly to the exogenous one, contributes to the inactivation of the epidermal gene program necessary for pluripotency establishment. It remains unclear however why the up-regulation of pluripotency genes, which are also targets of both KLF4 and HIC2, are not suppressed by *Hic2* overexpression. For example, KLF4 and exogenous HIC2 bind at the *Sall4* and *Nanog* loci at day 2 of reprogramming, but up-regulation of these genes are rather enhanced in *Hic2* overexpressing reprogramming (Supplementary Figure 3G and 3H). However, their accelerated up-regulation is observed only after day 4 or day 6 of reprogramming, respectively (Figure 3F and Supplementary Figure 3H). Hence, we speculate that the initial HIC2/KLF4 binding at the pluripotency gene loci does not directly affect their gene expression status and their accelerated up-regulation is an indirect effect. Alternatively, it is also possible that HIC2 directly contributes to the activation of pluripotency genes by recruiting co-activators that only become available later during reprogramming as HIC2 has also been reported to act as a transcriptional activator at least for one gene^39^. It could be interesting to investigate this possibility further in the future.

In this work, we demonstrated that the suppression of side effect gene activation by master TF overexpression can result in more direct and efficient cell conversion. Thanks to single-cell transcriptomics, it has become possible to investigate alternative cell identities, cell states and by-products generated during reprogramming in addition to the desired cell types in detail^12,30^. Identification and overexpression of suppressors that can prevent the generation of undesired cell states could render cell conversions more efficient and faithful. Overall, our study underscores the importance of understanding the molecular mechanisms of reprogramming processes to extend our capacity to manipulate cellular identity for use in medicine.

## Methods

### Cell culture

MEFs were cultured in MEF medium (Glasgow minimum essential medium [GMEM] supplemented with 10% fetal calf serum [FCS], 100 U/ml penicillin-streptomycin, 1x non-essential amino acids (Invitrogen), 1 mM sodium pyruvate, 2 mM glutamine, 0.05 mM 2-mercaptoethanol (Life Technologies) supplemented with 5 ng/ml fibroblast growth factor-2 (FGF2) and 1 ng/ml heparin). ESCs were cultured in ESC medium (MEF medium without FGF2 and heparin, supplemented with human LIF [leukaemia inhibitory factor], 100 U/ml), as described previously^34^. Reprogramming was performed in a reprogramming medium (ESC medium supplemented with 300 ng/ml (for transgenic MEF reprogramming) or 1ug/ml of doxycycline (for *piggyBac* reprogramming) (Sigma) and 10 µg/ml of l-ascorbic acid or 2-Phospho-L-ascorbic acid trisodium salt (Vitamin C) (Sigma). NSCs were cultured in NSC complete medium consisting of DMEM/F-12 Media, 1:1 Nutrient Mixture (Sigma), 1X N2 supplement (Thermo Fisher Scientific), 1X B27 supplement (Thermo Fisher Scientific), 8 mM glucose (Sigma), 100 U/ml Penicillin/Streptomycin (ThermoFisher Scientific), 0.001% Bovine Serum Albumin (BSA) (ThermoScientific), 0.05 mM β-mercaptoethanol (Thermo Fisher Scientific), supplemented with 10 ng/ml mouse Epidermal growth factor (EGF) (Peprotech) and 10 ng/ml human FGF2 (Peprotech).

### Plasmids

Plasmids and gRNAs used in this work are summarized in Supplementary Table S3. The plasmids and their sequences are available upon request.

### Lentivirus production and titration

1×10^6^ million HEK293T cells were cultured for 24 h in GMEM medium supplemented with 10% fetal calf serum (FCS), 1 mM glutamine and 100 U/ml Penicillin/Streptomycin (ThermoFisher Scientific) at 37 °C and 5% CO_2_. The following day, cells were transfected with a cocktail of plasmids containing 6 μg expression plasmid, 4.5 μg psPAX2 packaging vector, and 1.5 μg pMD2.G envelope vector mixed in 500 ul of 2X Hepes Buffered Saline (0.28 M NaCl, 0.05 M HEPES, 1.5 mM Na_2_HPO_4_, pH 7.05) to which 500 ul of 2.5 M CaCl_2_ solution is added drop-wise under constant agitation. The following day, the media was changed to GMEM and viral supernatant was collected 24 and 48h later and pooled, then cleared from cellular debris by centrifugation and filtered through a 0.45-μm syringe filter (Merck Millipore). The supernatant containing the gRNA virus particles was aliquoted and stored at –80 °C. For all other lentiviruses, viral supernatant was concentrated with Lenti-X™ Concentrator (Takara) according to the manufacturer’s guidelines. The concentrated lentiviruses were subsequently aliquoted and stored at –80 °C. For each experiment, lentiviruses were titrated on target cells either by flow cytometry or by immunofluorescence.

### ESC clonal assay with essential gene KO

*Cas9 TNG MKOS* ESCs were seeded at a density of 1×10^5^ cells per well in a 12-well gelatine-coated plate ^19^. On the following day, the cells were transduced at a multiplicity of infection (MOI) of 5 with sgRNA lentiviral particles containing 8 μg/ml polybrene (Millipore) for 6-8 hours, after which cells were given fresh ESC medium. ESCs were harvested 2 days later and BFP^+^ (blue fluorescent protein reporter contained in sgRNA carrying vectors) cells were flow-sorted with FACS AriaII (BD Biosciences) and plated at a clonal density (700 cells per well) in a 6-well gelatine-coated plate. The medium was replenished every other day. On day 11, cells were fixed with 4% paraformaldehyde for 10 minutes and washed with PBS. Whole-well imaging and quantification of Nanog-GFP^+^ colony numbers were performed with the Celigo S Cell Cytometer (Nexcelom).

### CFSE MEF proliferation assay

*Cas9 TNG MKOS* MEFs were seeded at a density of 8×10^4^ cells per well in a 12-well plate and transduced at MOI3 with lentiviral particles containing 8 μg/ml polybrene (Millipore) for 5 hours on the following day. The medium was then changed to MEF medium and cells were cultured for 2 days. MEFs were harvested and stained with 2.5 µM of CellTrace™ Far Red dye (Thermofisher Scientific) in PBS for 20 min at 37 °C and thoroughly washed with MEF medium. After staining, cells were plated at a density of 3.5×10^4^ cells per well in a 6-well plate and maintained in culture for 9 days. CellTrace™ Far Red-labelled cells at day 0 and day 9 were harvested and washed with FACS buffer (2% FCS in PBS) before acquisition with LSR Fortessa (BD Biosciences) cytometer. Dead cells were excluded using Propidium iodide staining (Sigma, 1 μg/ml). Data were analysed using Flowjo v10 proliferation platform and the division index (average number of divisions) was calculated based on the dilution of the fluorescent signals against day 0.

### Cas9 TNG MKOS transgenic MEF reprogramming

A total of 1 × 10^4^ *Cas9 TNG MKOS* MEFs were mixed with 9 × 10^4^ WT MEFs (129 strain) and seeded in gelatine-coated wells of 6-well plates. Cells were transduced with sgRNA lentiviruses at an MOI of 3 with 8 µg/ml polybrene (Merck-Millipore) for 4-6 h and then reprogramming was initiated by the addition of a reprogramming medium. On day 14, whole well colony images were taken using the Celigo S Cell Cytometer (Nexcelom) and the number of *Nanog*-GFP^+^ colonies was counted.

### *piggyBac* reprogramming of MEFs with overexpression of genes of interest

*Nanog*-GFP MEFs or wild-type MEFs isolated from E12.5 embryos were plated at 1-1.5 × 10^5^ cells per well in a gelatin-coated 6-well plate. The following day, co-transfection of a Dox-inducible *piggyBac* transposon vector carrying the *tetO-MKOS-ires-mOrange*, *tetO-STEMCCA-ires-mOrange* or *tetO-STEMCCA^+^*^9^*-ires-mOrange* cassette, *PB-CA-rtTA* vector carrying a *P2A*-linked to cDNAs of genes of interest, and *pCMV-hyPBase* was performed using 750 ng each DNA and 9 μl of FugeneHD (Promega) as per manufacturer’s instructions ^26,34,40^. 24 h later reprogramming was initiated with the reprogramming medium. The medium was changed every 2 days. In the case of wild type MEFs, cells were fixed with 4% paraformaldehyde for 10 minutes, permeabilized in 0.1% Triton-X in PBS for 2 hours, blocked in 5% BSA in PBS with 0.1% Tween20 for 1 hour at room temperature, and then stained in blocking solution with primary antibody for NANOG (eBioMLC-51, Thermofisher Scientific) overnight at 4 °C. The next day, an AlexaFluor488 conjugated secondary antibody (A-21208, Invitrogen) was applied in a blocking solution for 45 minutes at room temperature. For colony counting, whole well colony images were taken on day 14 using the Celigo S Cell Cytometer (Nexcelom) and colonies were counted with ImageJ.

### *piggyBac* reprogramming of NSCs with overexpression of genes of interest

NSCs were reprogrammed by nucleofection of a Dox-inducible *piggyBac* transposon vector carrying the *tetO-MKOS-ires-mOrange* cassette, *PB-CA-rtTA* vector carrying a *P2A*-linked to cDNAs of genes of interest and *pCMV-hyPBase* as previously described ^19^. 2 × 10^5^ NSCs were nucleofected with 750 ng each of the above-mentioned plasmids using SG Cell Line 4D Nucleofector X Kit (Lonza) and DN-100 program as per manufacturer’s instructions. Cells were recovered in NSC medium and then plated on a layer of wild-type MEF feeder cells seeded the day before at a density of 1 × 10^5^ cells per well in a gelatin-coated 6-well plate. One day post-nucleofection, reprogramming was initiated with NSC complete medium supplemented with 100 U/ml human LIF, 0.3 µg/ml of doxycycline (Sigma) and 10 µg/ml of L-ascorbic acid or 2-Phospho-L-ascorbic acid trisodium salt (Sigma). After 6 days, the medium was switched to serum-free N2B27-based medium (containing DMEM/F12 medium supplemented with N2 combined 1:1 with Neurobasal® medium supplemented with B27; all from Thermo Fisher Scientific), MEK inhibitor (PD0325901, 0.8 μM, Axon Medchem), GSK3b inhibitor (CHIR99021, 3.3 μM, Axon Medchem), 1 µg/ml of doxycycline (Sigma) and 10 µg/ml of L-ascorbic acid or 2-Phospho-L-ascorbic acid trisodium salt (Sigma). At day 16 of reprogramming, immunofluorescence for NANOG was performed as described for MEFs. Whole well images were taken using the Celigo S Cell Cytometer (Nexcelom) and colonies were counted with ImageJ.

### CD44, ICAM1, *Nanog*-GFP expression analysis during reprogramming

Cells harvested at different time points of reprogramming were stained in FACS buffer for 30 min at 4 °C and washed with FACS buffer (1% FCS in PBS) before acquisition with LSR Fortessa (BD Biosciences) cytometer. The following antibodies from eBioscience were used: ICAM1-biotin (13-0541-82; Dilution: 1/100), CD44-APC (17-0441-82; Dilution 1/300), streptavidin-PE-Cy7 (25-4317-82; Dilution: 1/1500). Dead cells were excluded using LIVE/DEAD™ Fixable Near-IR Dead Cell Stain Kit (ThermoFisher Scientific, Dilution: 1/1500). Data were analyzed using Flowjo v10.

### qPCR of epidermal genes

1 × 10^5^ MEFs were co-transduced with and rtTA lentiviruses as well as dox-inducible lentiviruses carrying either Ty1-BFP (control, BFP tagged with a Ty1 tag at the N-term end), *Ty1-Hic2* (HIC2 tagged with a Ty1 tag at the N-term end), or KLF4 at an MOI of 3 with 8 µg/ml polybrene (Merck-Millipore) for 4 h. After 2 days of culture in MEF media, 1 μg/ml of doxycycline (Sigma) was added to the refreshed MEF media. 96hrs later, total RNA was isolated using RNeasy Mini Kit (QIAGEN) according to manufacture’s instructions. Double strand complementary DNA (cDNA) was synthesized using SuperScript™ VILO™ cDNA Synthesis Kit (ThermoFisher Scientific). Real-time Polymerase Chain Reaction (PCR) was performed using Roche LightCycler 480 II Real-Time PCR System. Relative gene-expression levels were calculated by 2ΔΔCt method. Primers are listed in Supplementary Table S3.

## RNA-Seq

### Sample Preparation

For control and *Hic2* overexpressing MEF samples, 1 × 10^5^ *Cas9 TNG MKOS* MEFs were transduced with either *Ty1-BFP* (control, BFP tagged with a Ty1 tag at the N-term end) or *Ty1-Hic2* (HIC2 tagged with a Ty1 tag at the N-term end) lentiviruses at an MOI of 3 with 8 µg/ml polybrene (Merck-Millipore) for 4 h. After 3 days in culture in MEF media, the cells were harvested and FACS-sorted to exclude dead cells with LIVE/DEAD™ Fixable Near-IR Dead Cell Stain Kit (ThermoFisher Scientific, Dilution: 1/1500). For reprogramming samples, 0.5 × 10^4^ Cas9 *TNG MKOS* MEFs were mixed with 9.5 × 10^4^ WT MEFs (129 strain) and seeded in gelatin-coated wells of 6-well plates. Cells were transduced with either *Ty1*-*BFP* or *Ty1-Hic2* lentivirus at an MOI of 3 with 8 µg/ml polybrene (Merck-Millipore) for 4h, before being recovered for 24 h in MEF media. Reprogramming was initiated by the addition of a reprogramming medium. Cells were harvested at day 2, day 4, day 6 and day 10 of reprogramming, respectively, and a maximum of 1 × 10^5^ of mOrange^+^ OSKM and *Nanog*-GFP^+^ expressing cells were sorted with the FACS AriaII (BD Biosciences) per sample. Additionally, iPSCs were generated from *Cas9 TNG MKOS* MEFs transduced with a tet-inducible version of the Ty1-BFP and Ty1-Hic2 lentiviral particles *Nanog*-GFP^+^ iPSCs were harvested at day 15, and sorted with the FACS AriaII (BD Biosciences) and put in culture in ESC medium for 20 days in absence of doxycycline. All cells were homogenized with the QIAshredder kit (Qiagen) and total RNA was extracted from all samples using the RNeasy Plus Micro Kit (Qiagen). Libraries were prepped with the NEB Ultra II stranded mRNA Library prep kit (NEB). RNA-Seq libraries were sequenced with NextSeq, 75SE.

### Read processing

For each sequencing run, a quality control report was generated using FastQC and Illumina TruSeq adapter sequences were removed using Cutadapt^41^. Sequencing runs from the same biological sample were then concatenated and mapped to the GRCm38 reference genome using STAR^42^.

### Differential analysis

For each biological sample, aligned sequencing reads were first assigned to genomic features (e.g., genes) using Rsubread^43^ and a count table was generated. Differential expression analysis was then performed with DESeq2^44^, and statistically significant genes (e.g., FDR < 0.05 and log2FoldChange > 1) were identified using the standard workflow. Importantly, although the data represents a control and treatment time-series experiment, we opted to combine the factors of interest into a single factor for easier comprehension. GSEA was performed with GSEA (v.4.1.0) using the GSEAPreranked tool, whereby genes were pre-ranked based on their *P* values and fold changes.

### Downstream analysis

For exploratory analysis and visualization, a batch-corrected and regularized log matrix of expression values was used. The count table was first transformed to stabilize the variance across the mean using the rlog function from DESeq2 and then unwanted batch effects (e.g., library preparation date) were removed using the removeBatchEffect function from limma^45^. Gene ontology enrichment analysis was performed using the R package clusterProfiler (v 4.7.1.003) and visualized with enrichplot (v1.18.3). In Fig. 4H, the heatmap was generated using the pheatmap (v1.0.12) and LOESS (locally estimated scatterplot smoothing) curves were produced using the stat_smooth function of the ggplot2 (v3.4.2).

### ChIP-Seq

#### Sample Preparation

*Cas9 TNG MKOS* MEFs (3.5 × 10^6^ cells) were seeded onto 15-cm dishes (2 dishes in total) and cultured for 24h in MEF medium before infection with either a *Tet-On Ty1-BFP* or *Ty1-Hic2* lentiviruses at an MOI of 5. 48h later, cells were split across 5×15-cm dishes and media was changed to reprogramming the following day for a duration of 48h. For *Ty1-HIC2* (Ty1 tag is linked in the C-term of HIC2 for ChIP experiments) ChIP, cells were crosslinked with 2mM of Disuccinimidyl Glutarate (DSG) for 45 min at room temperature (RT) under constant agitation, before being cross-linked with 1% formaldehyde solution for 10 min RT. For KLF4 ChIPs, cells were only cross-linked with 1% formaldehyde solution for 10 min RT. Crosslinking was quenched by adding 120 mM glycine and incubating for 10 min at room temperature. Cells were collected using a scraper and pelleted by centrifugation at 1,350 × *g* for 5 min at 4 °C. The cross-linked pellets were washed 3 times with 10 ml of ice-cold PBS, and 3 to 4 aliquots of fixed cell pellets were made before flash freezing in liquid nitrogen and stored at –80 °C. Before nuclear extraction, 1 aliquot of cell pellets per ChIP was thawed on ice for 2–3 h and suspended in 10 ml filtered, ice-cold lysis buffer (50 mM HEPES-KOH pH 7.5, 140 mM NaCl, 1 mM EDTA, 10% glycerol, 0.5% NP-40, 0.25% Triton X-100 and 1 tablet of complete ultra-protease inhibitor cocktail (Roche)). The suspension was gently mixed on a rotating platform at 4 °C for 10 min. The cells were disrupted using a 7 ml glass-Dounce homogenizer (40 strokes) on ice. The nuclei were pelleted by centrifugation (1,350 × *g* for 5 min at 4 °C) and washed with 10 ml ice-cold wash buffer (10 mM Tris-HCl pH 8, 200 mM NaCl, 1 mM EDTA, 0.5 mM EGTA and 1 tablet of complete ultra-protease inhibitor cocktail (Roche)) for 10 min at 4 °C. The nuclei were collected by centrifugation and resuspended in 4 ml sonication buffer (10 mM Tis-HCl pH 8, 100 mM NaCl, 1 mM EDTA, 0.5 mM EGTA, 0.1% sodium-deoxycholate, 0.5% *N*-lauroylsarcosine and 1 tablet of complete ultra-protease inhibitor cocktail (Roche)). The resuspended nuclei were split into five aliquots in pre-chilled 1 ml milli-tubes containing AFA Fibre and sonicated using a Covaris-M220 focused-ultrasonicator (Covaris). Each milli-tube was sonicated for 10 min and kept on ice for 30 min per sonication cycle for a total of 4 cycles. Sonicated chromatin was transferred to Protein-Lobind tubes (Eppendorf). A total of 100 µl of 10% Triton X-100 was then added to each 1 ml sonicated chromatin to increase solubility. Chromatin samples were then centrifuged (20,000 × *g* at 4 °C for 10 min) and the supernatants were pooled into fresh tubes. A 50-µl aliquot of each pooled sample was analysed to check the fragment size distribution and to quantify the DNA content of the resulting sonicated chromatin using a Nanodrop spectrophotometer. Another 50-µl aliquot was retained to be used as input for ChIP analysis. The sonicated chromatin and the input sample were snap-frozen in liquid nitrogen and stored at –80 °C.

For each ChIP analysis, 30 µl Dyna Protein-G magnetic beads (Thermo Fisher Scientific) were washed three times with 1 ml blocking solution (0.5% w/v BSA in PBS/Tween-20). The beads were saturated with the appropriate antibody by adding 10 µg of Ty1 (Diagenode) or KLF4 (R&D) antibody and incubated on a rotating platform at 4 °C for at least 6 hours. Sonicated chromatin (60 µg) was then mixed with the antibody-saturated beads and incubated on a rotating platform at 4 °C overnight. The beads were transferred to a fresh tube, washed 5 times with 1 ml wash buffer (50 mM HEPES-KOH pH 7.6, 500 mM LiCl, 1 mM EDTA, 1% NP-40 and 0.7% sodium-deoxycholate) and then washed once more with TE buffer (10 mM Tris-HCl pH 8, and 1 mM EDTA) containing 50 mM NaCl. The bound chromatin was eluted by incubating the beads in 210 µl EB (50 mM Tris-HCl pH 8, 10 mM EDTA and 1% SDS) at 65 °C for 30 min. The beads were pelleted by centrifugation at 16,000 × *g* for 1 min, and 200 µl supernatant containing the soluble chromatin was transferred to fresh tubes. The crosslinking was reversed by incubating the eluted chromatin and input DNA (in EB) at 65 °C for 16 h with shaking. An equal volume of TE (200 µl) was added to the eluted chromatin and input DNA to reduce the concentration of SDS. RNA in the samples was digested by adding 0.2 mg/ml RNase-A for 2 hours at 37 °C. Proteins were then digested by adding 0.2 mg/ml proteinase-K for 2 hours at 55 °C. ChIP DNA and input DNA were subsequently purified by phenol-chloroform extraction followed by ethanol precipitation and resuspended in EB buffer (100 mM Tris pH 8.0) before DNA concentrations were measured by Qubit Fluorometric Quantitation (Invitrogen, Thermo Fisher Scientific). DNA from ChIP assays and inputs were prepared using the NEBNext Ultra-II DNA Library Prep kit (New England Biolabs) according to the manufacturer’s instructions. Each library was uniquely barcoded using NEBNext Multiplex Oligos for Illumina (Dual-Index Primers Set-1) (New England Biolabs). The quality of the DNA libraries was assessed using Agilent HS-DNA-Screen Tape (Agilent). Samples were sequenced with the NextSeq High 40PE machine (Illumina).

### Data analysis

PCR duplicates were removed using the Picard MarkDuplicates (v2.23.4) (http://broadinstitute.github.io/picard/). Uninformative alignments were subsequently filtered using a combination of SAMtools (v1.10) and BEDtools (v2.29.2) commands^46,47^. Specifically, reads mapped to the mitochondrial chromosome and blacklisted regions were filtered. For KLF4 ChIPs, BAM files from two biological replicates were merged using SAMtools. Peak calling was performed with MACS2 (v2.2.6) with subtraction of input background^48^. For Ty1 ChIPs, Ty1-immunoprecipitated DNA from Ty-BFP overexpressing cells at 48h of MKOS expression was used as background. The correlation plot of read coverage was produced using the multiBigwigSummary and plotCorrelation commands from deepTools (v3.3.2)^49^. Heatmaps of read coverage were produced using the computeMatrix and plotHeatmap commands from deepTools. Motif enrichment analysis was performed using the MEME-ChIP tool from the MEME suite (v5.1.1)^50^. OCT4, SOX2 and KLF4 binding data in ESCs are taken from GSE90893^51^

## scRNA-Seq

### Sample Preparation

*Nanog*-GFP MEFs were reprogrammed using the *piggyBac MKOS* reprogramming system with overexpression of genes of interest (*BFP* as a control, or *Hic2*) described earlier. Cells were collected on days 2, 4, 6 9 and 12, alongside *Nanog*-GFP MEFs and *Nanog*-GFP ESCs. 2×10^6^ cells per sample were labelled with a unique Cell Multiplexing Oligo (CMO) provided in the 10x Genomics 3’ CellPlex Kit according to the manufacturer’s instructions. Subsequently, cells were sorted with the FACS AriaII (BD Biosciences) for either mOrange (reporting for MKOS expression), GFP (reporting for NANOG expression, or both mOrange and BFP. Non-viable cells were excluded based on DRAQ7. In the case of MEFs, cells were only sorted based on DRAQ7 signal. After confirming the viability of cells after sorting using trypan blue and a haemocytometer, 10000 single cells per every 6 samples were pooled together, generating 2 pools that were each processed through the Chromium Single Cell Platform using a Chromium Next GEM Single Cell 3′ GEM Library and Gel Bead kit (v.3.1 chemistry, 10x Genomics) and a Chromium Next GEM Chip G kit following the manufacturer’s instructions. Libraries were sequenced using NextSeq 2000 platform (Illumina).

### Data analysis

The sequenced libraries were demultiplexed and aligned to mm10 reference genome using the multi-function from Cell Ranger (10x Genomics, v6.1.2) with the parameter min-assignment-confidence 0.8 and the cmo-set option including only the used tags as the CMO reference. The gene count matrices were processed using the R package Seurat (v4.1.0)^31^. Cells expressing less than 200 genes, over 150,000 UMI counts, and more than 10% mitochondrial counts were filtered. The filtered data were then log-normalized and scaled, and data from different libraries were merged using the FindIntegrationAnchors and IntegrateData functions of Seurat. Followed by dimensionality reductions using the RunPCA and RunUMAP functions, clustering was performed using the FindNeighbours function with the top 30 principal components and the FindClusters function with the resolution = 0.4. The force-directed layout (FDL) graph was generated using the scanpy.tl.draw_graph function of scanpy (v1.7.3) with the UMAP coordinates for initialization^52^. The shiny app for interactive analysis (https://kkaji.shinyapps.io/230420shiny/?_ga=2.71125584.1988667254.1682608652-2076402475.1682608652) was generated using the R package ShinyCell (v2.1.0)^53^. The scRNA-seq data of the reprogramming process reported by Schiebinger, et al. was downloaded from GSE122662^15^. The metadata was obtained from https://broadinstitute.github.io/wot/. Since this data contained ∼250K cells, we randomly selected 2,000 cells each from the cells classified as MET, stromal, epithelial, neural, trophoblast, and iPSC. This selected dataset (12,000 cells) and our scRNA-seq data (13,315 cells) were each normalized and scaled using the SCTransform function and regressing out the UMI count with the “glmGamPoi” method^54^, and integrated using the PrepSCTIntegration, FindIntegrationAnchors, and IntegrateData functions of Seurat. Dimensionality reduction and visualization were performed as described above.

### Data availability

Raw and processed bulk RNA-seq, scRNA-seq, ChIP-seq data are available at ArrayExpress from the following links, respectively.

https://www.ebi.ac.uk/biostudies/arrayexpress/studies/E-MTAB-13169?key=b515b122-a4e2-4fca-9aa8-58df8c9c2be7

https://www.ebi.ac.uk/biostudies/arrayexpress/studies/E-MTAB-13029?key=fc47d3b3-2fed-4241-92ef-9b071a873d19

https://www.ebi.ac.uk/biostudies/arrayexpress/studies/E-MTAB-13170?key=a5b4a609-e1a1-49dd-9051-1853044328b3

## END NOTE

## Acknowledgements

We thank I. Chambers for providing the TNG ESC line, F. Rossi and C. Cryer for assistance with flow cytometry, Biomed unit staff for mouse husbandry, EMBL GeneCore for RNA-seq, A. Corsinotti for scRNA-seq, A. Soufi, and M.L. Huynh for comments on the manuscript. Some of the computations for this work were enabled by resources provided by the Swedish National Infrastructure for Computing (SNIC) at the Uppsala Multidisciplinary Center for Advanced Computational Science (UPPMAX) partially funded by the Swedish Research Council through grant agreement no. 2018-05973. This work was supported by MRC senior non-clinical fellowship (MR/N008715/1) funded for K.K. K.Y. was supported by the Wellcome Trust (206194). D.F.K., J.A. and M.Y. were supported by the BBSRC (EASTBIO doctoral training partnership), Principal’s Career Development scholarship from the University of Edinburgh, and Japan Society for the Promotion of Science (JSPS) Overseas Research Fellowships, respectively. V.O. and E.A. were supported by the Swedish Foundation for Strategic Research (A3 04 159p). V.O. was also supported by the Swedish Research Council (Vr 621-2008-3074). S.K. was supported by Jane and Aatos Erkko Foundation. RNA-sequence was performed in EMBL GeneCore Facility.

## Author Contribution

M.B. designed and performed experiments. M.Y., J.A. and S.K. contributed to the analyses of the RNA-seq, scRNA-seq and ChIP-seq data sets. D.F.K. has performed the CRISPR/Cas9 KO screen and S.Z., M.O. and A.S. provided technical support. E.A. and V.O. generated the screening data website. K.Y. provided the reagents for CRISPR/Cas9 KO experiments and advised on the original screen. K.K. conceived the study, supervised experiment design and data interpretation, and wrote the manuscript with M.B.

---

**Supplementary Figure 1.**
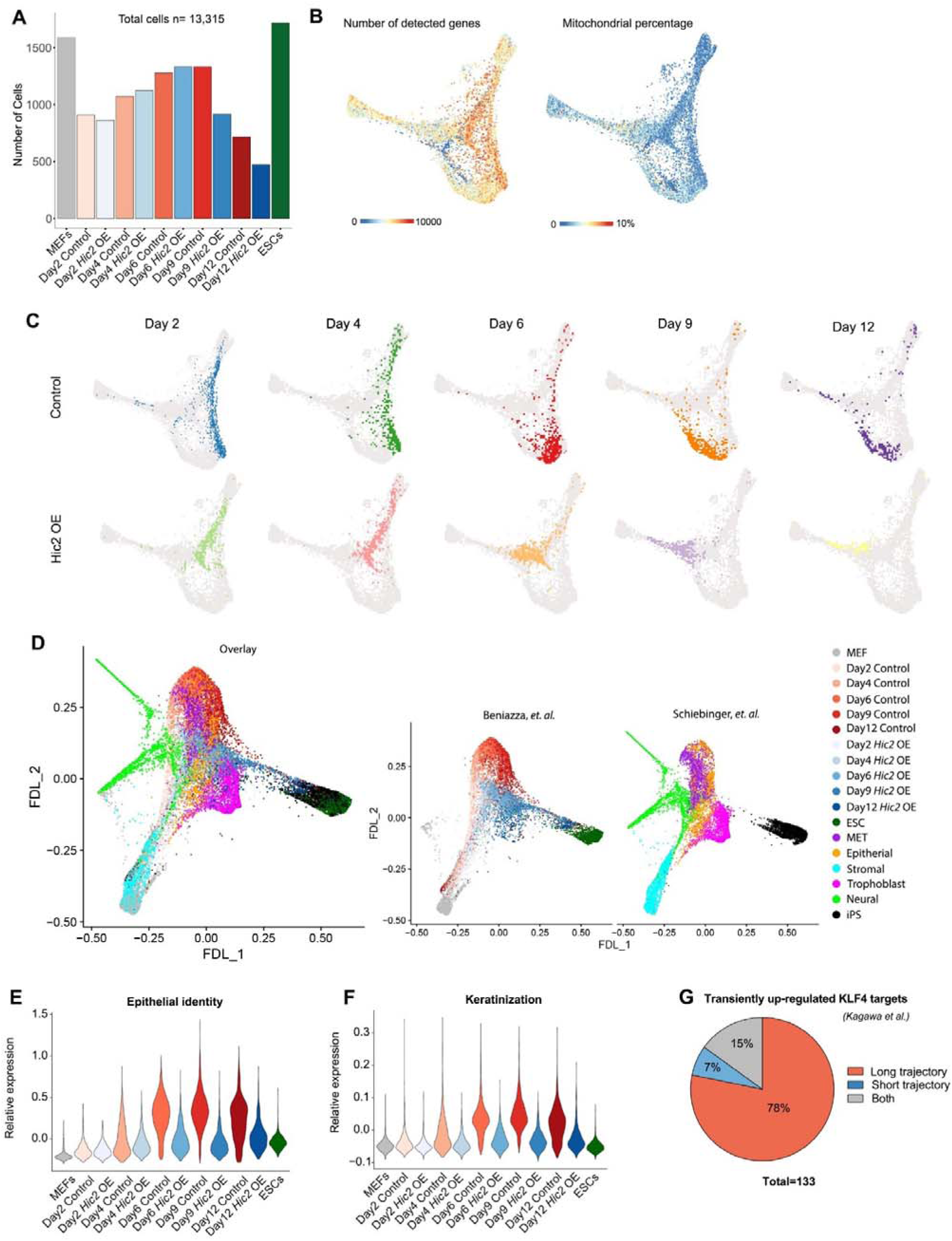
scRNA-seq during reprogramming with and without *Hic2* overexpression. **A**. The number of cells with high-quality sequence data from different samples. **B**. Number of detected genes per cell and proportions of mitochondrial genes. **C.** Day-by-day transition of cells during reprogramming with (bottom) and without (top) *Hic2* overexpression (OE) on the FDL plot. **D.** Overlay with scRNA-seq data from OKSM secondary MEF reprogramming by Schiebinger, *et. al.,* using randomly selected 2,000 cells each from the cells classified as MET, stromal, epithelial, neural, trophoblast, and iPSC. **E.** Expression of epithelial gene signature^15^ in control and *Hic2* OE samples. **F.** Expression of genes within the “keratinization” gene ontology in control and *Hic2* OE samples. **G.** Proportions of known transiently up-regulated KLF4 targets^10^ within the long and short reprogramming trajectories

**Supplementary Figure 2.**
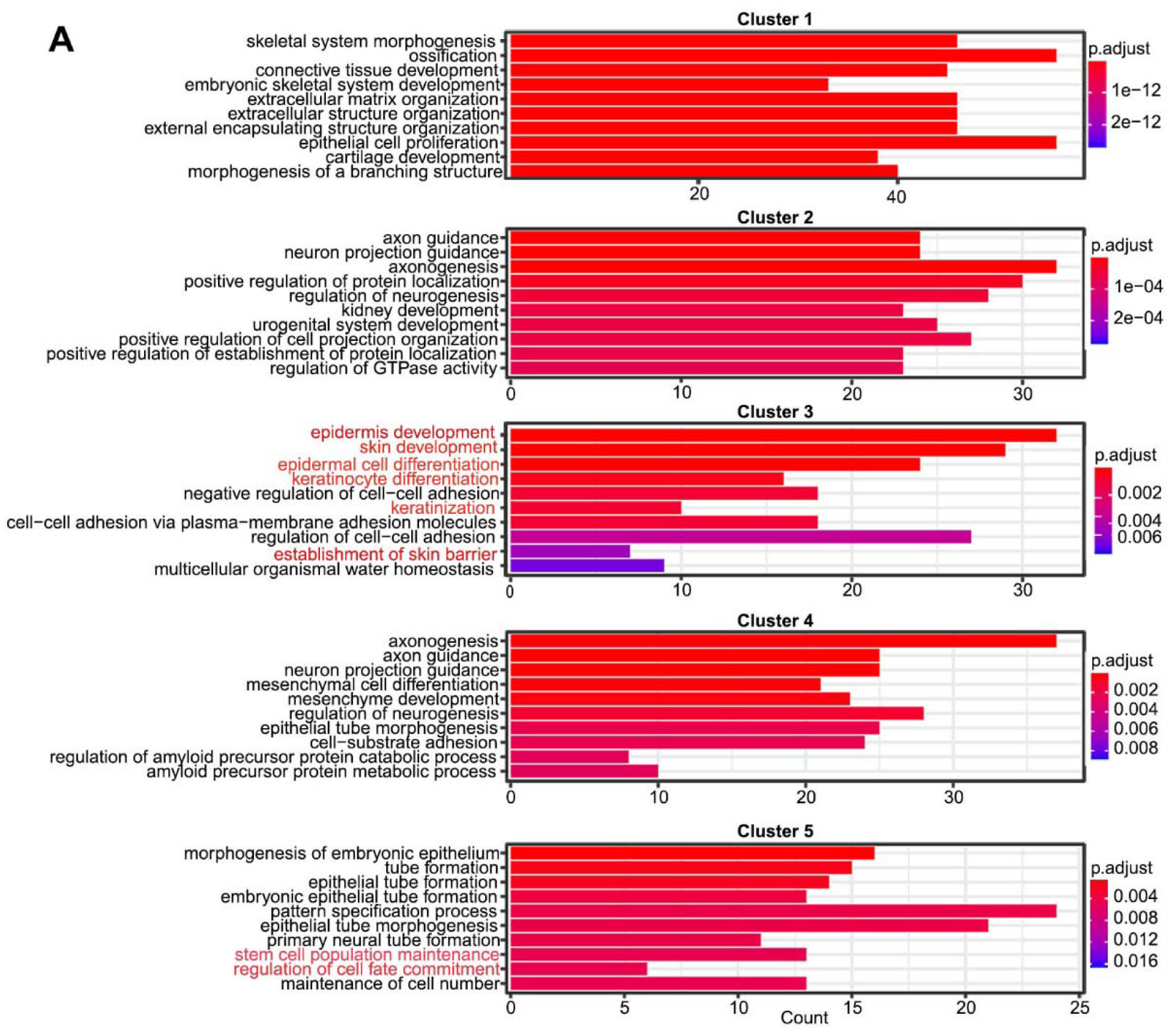
Top 10 enriched gene ontology terms in the clusters from Figure 4H.

**Supplementary Figure 3.**
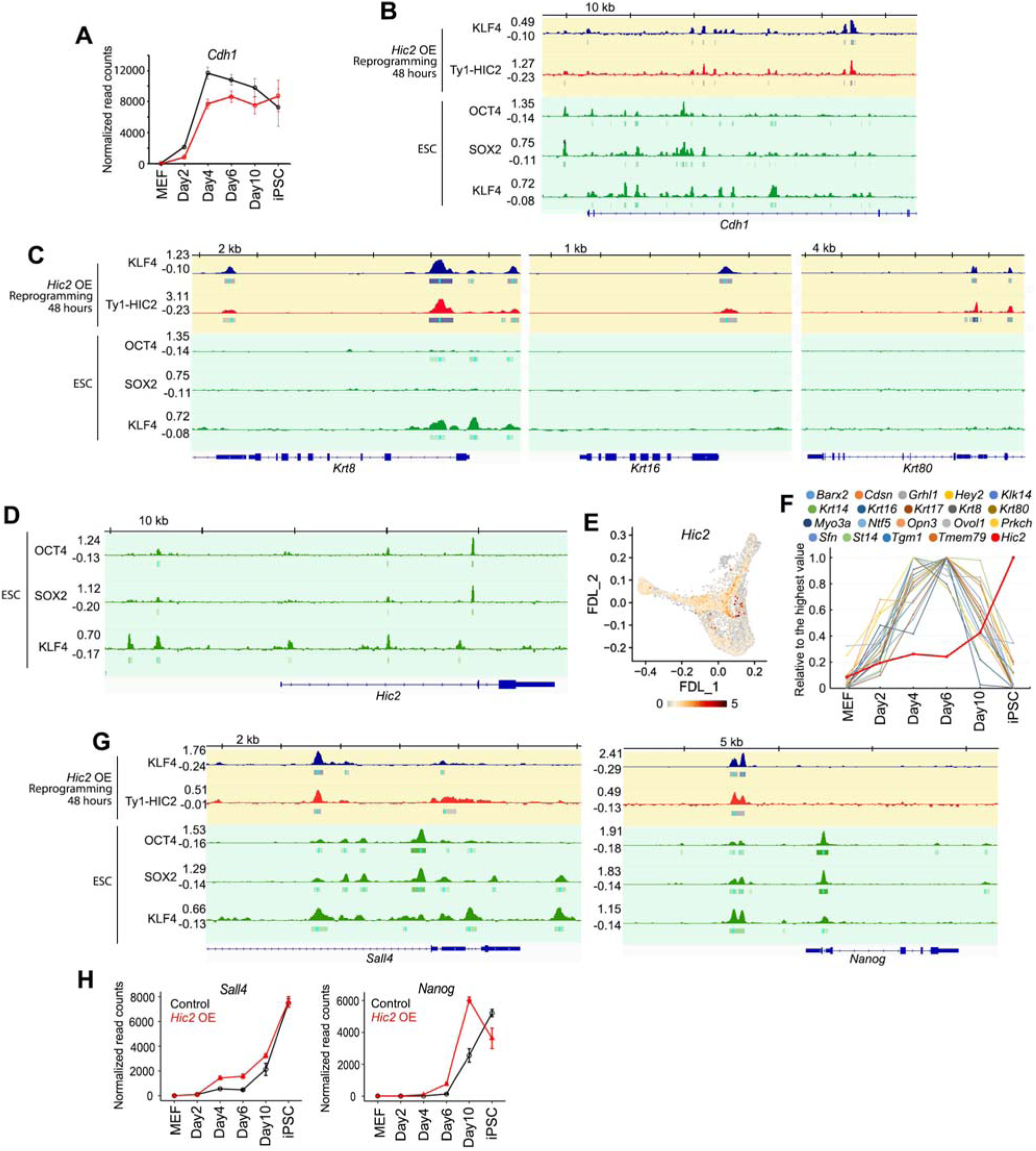
Potential factors involved in *Hic2, Nanog and Sall4* induction during reprogramming. **A.** Expression of *Cdh1* during reprogramming with and without *Hic2* OE. Error bars indicate SEM from 3 replicates. **B.** Binding of KLF4 and HIC2 at the *Cdh1* locus at 48 hours of reprogramming with *Hic2* OE and in ESCs (GSE90893). **C.** KLF4 and HIC2 binding at 48 hours of reprogramming with *Hic2* OE and in ESCs at 3 epidermis gene loci. **D.** OCT4, SOX2, KLF4 biding in the Hic2 locus in ESCs. **E.** *Hic2* expression during reprogramming with/without Hic2 overexpression (OE) **F.** Expression of epidermal/epithelial/keratinocyte genes from the bulk RNA-seq data of reprogramming without Hic2 overexpression. Data represent an average of 3 independent experiments, normalized to the highest values in each gene. **G.** KLF4 and HIC2 binding at 48 hours of reprogramming with *Hic2* OE and in ESCs at the *Nanog* and *Sall4* loci. **H.** Expression of *Nanog* and *Sall4* during reprogramming with and without *Hic2* OE. Error bars indicate SEM from 3 replicates.

